# Intrinsically disordered linkers determine the interplay between phase separation and gelation in multivalent proteins

**DOI:** 10.1101/164301

**Authors:** Tyler S. Harmon, Alex S. Holehouse, Michael K. Rosen, Rohit V. Pappu

## Abstract

Many intracellular membraneless bodies appear to form via reversible phase transitions of multivalent proteins. Two relevant types of phase transitions are *sol-gel transitions* (gelation) and *phase separation plus gelation*. Gelation refers to the formation of a system spanning molecular network. This can either be enabled by phase separation or it can occur independently. Despite relevance for the formation and selectivity of compositionally distinct protein and RNA assemblies, the determinants of gelation as opposed to phase separation plus gelation remain unclear. Here, we focus on linear multivalent proteins that consist of interaction domains that are connected by disordered linkers. Using results from computer simulations and theoretical analysis we show that the lengths and sequence-specific features of disordered linkers determine the coupling between phase separation and gelation. Thus, the precise nature of phase transitions for linear multivalent proteins should be biologically tunable through genetic encoding of or post-translational modifications to linker sequences.

## Introduction

There is growing interest in intracellular phase transitions that are thought to be important in the formation of membraneless organelles and other protein / RNA bodies, collectively known as biomolecular condensates (1). These are two- or three-dimensional assemblies that comprise of multiple proteins and RNA molecules and lack a surrounding membrane. Many biomolecular condensates are manifest as different types of membraneless organelles and other assemblies that are involved in cell signaling (2), ribosomal biogenesis (3-5), cytoskeletal regulation (6, 7), stress response (8-11), cell polarization (12, 13), and cytoplasmic branching (14). It has been proposed that the protein components of biomolecular condensates can be parsed into scaffolds and clients (15). Scaffolds are thought to drive phase transitions, whereas client molecules preferentially partition from the cytoplasm or nucleoplasm into the biomolecular condensates (15, 16). Scaffold proteins have distinct features, the most prominent being *multivalency* of well-folded protein domains or short linear motifs (SLiMs) that are encompassed in low complexity disordered regions (1, 6, 17-19). The concept of valence refers to the number of interaction domains or SLiMs within a multivalent protein. Ligands of multivalent proteins can be other multivalent proteins or polynucleotides. The simplest multivalent proteins are linear polymers that consist of multiple protein interaction domains or SLiMs that are connected to one another by intrinsically disordered linkers that may or may not lack specific interaction motifs (**Figure 1a**). These systems can undergo two types of reversible phase transitions, namely a solution-to-gel (sol-gel) transition (gelation) or phase separation plus gelation.

**Figure 1:**
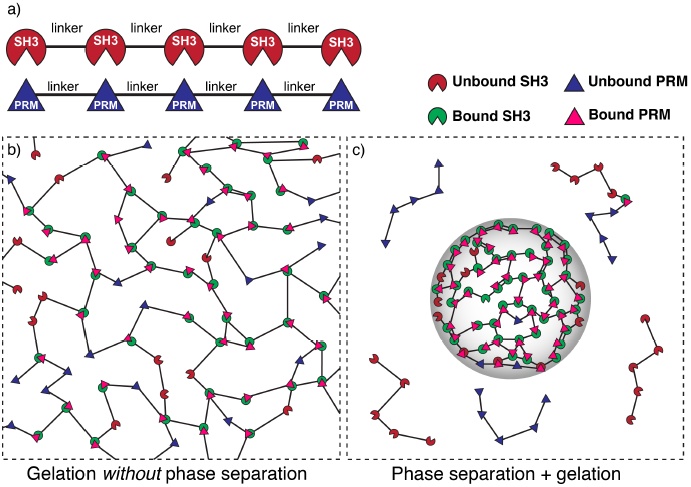
Depiction of gelation without phase separation as opposed to phase separation plus gelation. (a) Schematic of a synthetic multivalent system. SH3 domains bind to proline-rich modules (PRMs). Multivalent SH3 and PR proteins result from the tethering of multiple SH3 domains (or PRMs) by linkers. (b) Schematic of gelation without phase separation: If the bulk concentration of interaction domains is above the gel point but below the saturation concentration then a system spanning network forms across the entire system volume. In this scenario, a percolation transition is realized without phase separation. (c) Schematic of phase separation plus gelation. Linker-mediated cooperative interactions of multivalent proteins drive phase separation, depicted here as a confinement of molecules into a smaller volume (gray envelope) when compared to the system volume (dashed bounding box). If the bulk concentration of interaction domains is higher than a saturation concentration then a dense phase comprising of multivalent SH3 and PRM proteins will be in equilibrium with a dispersed phase of unbound proteins. A droplet-spanning network will form because the concentration of interaction domains within the dense phase is above the gel point.

*Gelation* refers to a switch from a solution of dispersed monomers and oligomers – a sol – to a system-spanning network – a gel (**Figure 1b**). This *connectivity transition* is characterized by the existence of a concentration threshold, known as the percolation threshold (20) that defines the *gel point* (21-23). If the bulk concentration of interaction domains is below the gel point, then the multivalent proteins form a sol. Above the gel point, the multivalent proteins and their ligands are incorporated into a system spanning network known as a gel. Here, we focus on physical gels (24), which are defined by specific, reversible non-covalent interactions, that represent physical crosslinks between protein modules / SLiMs and their ligands (2, 25). Our definition of a physical gel is based on Flory’s work (26) and is consistent with criteria outlined by Almdal et al. (27). *By these definitions, a gel is a percolated network characterized by system spanning physical crosslinks*. These definitions are agnostic about the material properties of gels. Specifically, gels are not automatically conflated with solids nor do we postulate that gels have to be pathological states of matter.

Polymer solutions can also undergo *phase separation* (17, 28-30). Given the three-way interplay among polymer-solvent, solvent-solvent, and polymer-polymer interactions, a necessary condition for phase separation is that inter-polymer attractions are more favorable, on average, than all other interactions (17, 24, 31). Above a *saturation concentration*, the polymer solution will undergo liquid-liquid phase separation (LLPS) by separating into a dense polymer-rich phase that coexists with dilute liquid, deficient in polymers (28, 29). The formation of two distinct phases characterized by LLPS represents a *density transition*, with the dense phases typically forming spherical droplets (**Figure 1c**).

Although phase separation and gelation involve changes to two distinct physical properties of the system, these transitions can be coupled to one another. Phase separation of multivalent proteins will always promote gelation if the concentration of interaction domains within the dense phase is above the gel point (**Figure 1c**). Therefore, there are clearly two distinct types of phase transitions to consider for multivalent proteins: *sol-gel transitions* (gelation) as opposed to *phase separation plus gelation*. Sol-gel transitions are continuous transitions. Prior to gelation there is a pure sol of monomers and oligomers. Gelation refers to the crossover in the extent of crosslinking that yields a percolated or system spanning network, *i.e*., a gel. In sol-gel transitions, sols and gels do not coexist. Instead, a sol changes continuously to a gel. Phase separation plus gelation is a discontinuous, first-order transition because the dilute phase, a sol, will coexist with the dense phase, a gel. In this scenario, the gel is actually a droplet-spanning network.

*The question of interest is what drives the extent and type of coupling between phase separation and gelation in multivalent proteins*? Using computer simulations and theories we show that the physical properties of linkers and the affinities between interaction domains are key determinants of the coupling between phase separation and gelation in linear multivalent proteins. Specifically, we show that for linear multivalent proteins of fixed binding-module affinity and valence, the disordered linkers determine the preference for phase separation plus gelation as opposed to gelation without phase separation. This behavior is determined by the sequence-specific properties of linkers, which can be quantified in terms of a single parameter known as the effective solvation volume.

*Effective solvation volumes are defined as the volumes associated with pairs of linker residues for interactions with the surrounding solvent as opposed to interactions with themselves* (31). The effective solvation volume (v_es_) of a linker can be pictured in terms of the impact a linker has on bringing together interaction modules that are connected to either end (see **Figure 2**). Qualitatively, we can think about this in terms of a hypothetical outwards force that acts on the two interaction modules at either end of the linker. When v_es_ is positive, the linker is highly expanded and this outwards force repels the two interaction modules, driving them apart. A positive v_es_ is realized because the linker is self-repelling, carving for itself a large volume in space for favorable interactions with the solvent. When v_es_ is negative, the linker is compact, and the hypothetical outwards force pulls the two interaction modules in, driving them close together. A negative v_es_ is realized because the solvent is squeezed out, the linker is self-attractive, and this causes the interaction domains to be pulled towards one-another. When v_es_ is close to zero, the linker does not have strong interaction preferences and mimics a passive tether. Accordingly, both expanded and compact linker conformations are equally likely. The hypothetical outwards / inward force is negligible – the preferences for compact versus expanded conformations cancel one another – and the interaction modules meander around in three-dimensional space with respect to one another, restrained only the connectivity of the linker. A value of v_es_ ≈; 0 is realized due to a counterbalancing of attractive and repulsive interactions in the linker.

**Figure 2:**
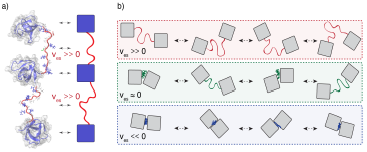
Illustration of the impact of linker effective solvation volumes on the conformational fluctuations and inter-domain distances in linear multivalent proteins. (a) Schematic of three SH3 domains connected by positive v_es_ linkers. In a cartoon schematic, the SH3 domains are shown as blue squares and the linkers are depicted as red tethers. The bidirectional arrows indicate the mapping between the molecular structures and the cartoon schematic. (b) Comparative schematics of SH3 domains connected by different types of linkers. The top row shows a pair of domains connected by linkers of high positive effective solvation volumes. For linkers with near zero effective solvation volumes, the inter-domain distances are characterized by large fluctuations and this engenders large concentration fluctuations. The bottom row shows the scenario for domains connected by linkers with negative v_es_ values. In this scenario, the inter-domain distances seldom exceed the sum of the individual radii of gyration.

The effective solvation volume of a linker can be quantified in terms of the solvent-mediated pairwise interactions between pairs of linker residues and the details are discussed in **Appendix A**. If the linker sequence is such that there are *net* attractions between all pairs of residues, then v_es_ will be negative and this will be true for linkers that form compact globules. Conversely, if there are *net* repulsions between all pairs of residues, then the residues prefer to be solvated and v_es_ will be positive. This is the case for so-called self-avoiding random coil (SARC) linkers. Finally, if the effects of inter-residue attractions offset the effects of inter-residue repulsions, then v_es_ ≈ 0 and this is the scenario for so-called Flory random coil (FRC) linkers. The effective solvation volume is directly proportional to the second virial coefficient denoted as *B*_2_ (31, 32). Negative, zero, or positive values of v_es_ correspondingly imply negative (attractive interactions), zero (non-interacting), or positive (repulsive interactions) values of *B*_2_. Therefore, v_es_ can be inferred using either atomistic simulations (as shown in this work) or via measurements of *B*_2_ as shown by Wei et al. (32).

For generic homopolymers, the sign and magnitude of v_es_ are determined by the effective chain-solvent interactions, which in turn depend on the chemical makeup of the chain. For proteins, the interplay between chain-chain and chain-solvent interactions is specified by the amino acid sequence, whereby the composition and patterning of a disordered linker will determine the balance of chain-chain and chain-solvent interactions (33-35). Therefore, the effective solvation volume of a disordered linker is determined directly by its primary sequence.

**Our work is designed to answer a simple question: How do changes to the effective solvation volumes and lengths of disordered linkers influence the coupling between phase separation and gelation for linear multivalent proteins?** To set the stage for our investigations, we first performed proteome-wide bioinformatics analysis combined with all-atom simulations to quantify conformational consequences of sequence-specific effective solvation volumes of disordered linkers in naturally occurring multi-domain human proteins. This analysis shows that the sub-proteome of linear multivalent proteins comprises of linkers of varying lengths that span a range of effective solvation volumes, from significantly negative to significantly positive values. Using coarse-grained numerical simulations and analytical theories we then show that the coupling between phase separation and gelation in linear multivalent proteins is directly determined by the physical properties of linkers, which include the lengths of linkers and their sequence-specific effective solvation volumes.

## Results

### Disordered linkers between folded domains in the human proteome span the entire range of effective solvation volumes

We first sought to obtain accurate and efficient estimates of the effective solvation volume (v_es_) for a large set of disordered segments. For this we used all-atom simulations, which have a proven track record of describing sequence-specific conformational properties of intrinsically disordered proteins (33, 35-37). Although a formal and rigorous calculation of v_es_ is technically possible using these simulations, this approach is computationally expensive and non-trivial for large numbers of sequences. Recognizing that the effective solvation volume directly determines the global dimensions of a linker, we used the ensemble-averaged conformational properties to calculate a proxy for v_es_ (38). Specifically, we leverage the profile of inter-residue distances to determine how a given linker sequence deviates from a sequence-specific theoretical reference that recapitulates v_es_ = 0, which is the Flory Random Coil (FRC) (39). These profiles (**Figure 3a**) describe the average spatial separation between all pairs of residues as a function of their separation along the polypeptide sequence.

We obtained sequence-specific inter-residue distance profiles by performing all-atom Metropolis Monte Carlo simulations using the ABSINTH implicit solvent model and forcefield paradigm (40) as described in the methods section. **Figure 3a** shows the calculated inter-residue distance profiles for fourteen distinct sequences, each of length 40 residues. Details of the sequences are shown in (**Table 1**). **Figure 3a** illustrates changes to the inter-residue distance profiles as a function of changes to the fraction of charged residues. **Figure 3a** also shows the inter-residue distance profile for a reference FRC linker. Sequences with positive v_es_ will have inter-residue distance profiles that lie above the FRC reference. Conversely, sequences with negative v_es_ will have profiles with uniformly smaller inter-residue spatial separations for given sequence separations when compared to the FRC reference. Accordingly, **Figure 3a** shows that sequences deficient in charged residues are expected to have negative v_es_ values, whereas sequences enriched in charges are expected to have positive v_es_ values.

**Table 1:**
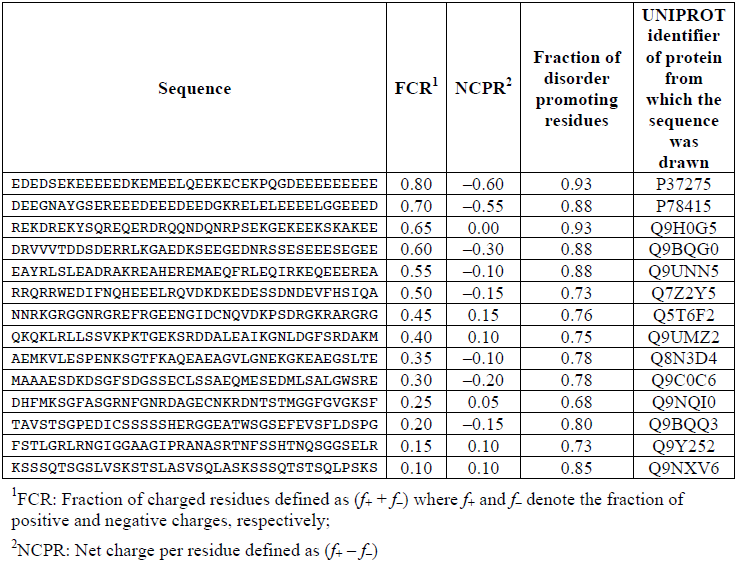
Details of the fourteen sequences chosen at random from the human proteome. All sequences have identical lengths (40 residues) and are enriched in disorder promoting residues. The sequences are listed in descending order of the fraction of charged residues. ^1^FCR: Fraction of charged residues defined as (*f*_+_ + *f*_–_) where *f*_+_ and *f*_–_ denote the fraction of positive and negative charges, respectively; ^2^NCPR: Net charge per residue defined as (*f*_+_ – *f*_–_)

**Figure 3:**
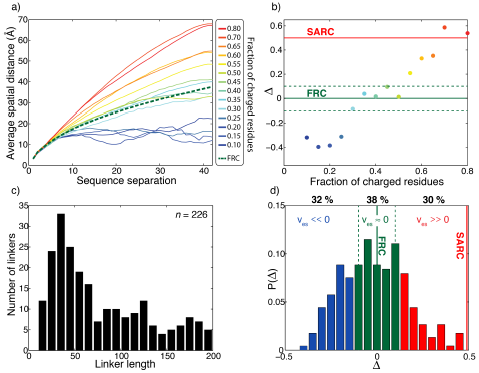
Effective solvation volumes for disordered linkers from the human proteome. (a) Inter-residue distance profiles for fourteen representative sequences, each 40-residues long. The legend shows the fraction of charged residues within each linker. The green dashed curve shows the inter-residue distance profile for the reference FRC limit. (b) Summary of the variation of Δ as a function of the fraction of charged residues for thefourteen representative sequences. Here 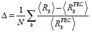 is the number of linker residues, ⟨*R*⟩ is the average spatial separation between residue pairs that are *k* apart in the linear sequence, 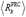 is the corresponding spatial separation for a FRC chain, and the summation index *k* runs across all sequence-separations. Linkers for which Δ < –0.1 will have negative effective solvation volumes (v_es_ < 0); linkers for which –0.1 ≤ Δ ≤ 0.1 will have near zero effective solvation volumes (v_es_ ≈ 0); and linkers for which Δ > 0.1, will have positive effective solvation volumes (v_es_ > 0). For the self-avoiding random coil (SARC) linkers, Δ ≈ 0.5 and this is shown as a horizontal red line. (c) Length distribution of all 226 unique disordered linkers. (d) Distribution of Δ values extracted from all-atom simulations of all 226 linkers. We delineate the Δ-distribution into three regimes: Δ < –0.1 (blue bars), –0.1 ≤ Δ ≤ 0.1 (green bars), and Δ > 0.1 (red bars). These regimes correspond, respectively to linkers for which v_es_ is less than zero, near zero, or greater than zero.

Since inter-residue distance profiles are direct manifestations of sequence-specific effective solvation volumes (38), we use these profiles to calculate a parameter Δ that serves as a proxy for estimating sequence-specific v_es_ values. This parameter is defined as the mean signed difference between the sequence-specific inter-residue distance profile and the corresponding profile for a FRC reference. In **Figure 3b** we plot the calculated Δ values against the fraction of charged residues for the fourteen disordered sequences from **Figure 3a**. The value of Δ can be negative, equal to zero, or positive and this depends on whether the value of v_es_ is negative, zero, or positive, respectively. Sequences that form compact globules have negative values of v_es_ and negative values of Δ. This is true for sequences with fractions of charged residues below 0.3. Within an interval between 0.3 and 0.5 for the fraction of charged residues, sequences mimic the FRC limit, where v_es_ ≈ 0. This is manifest as –0.1 ≤ Δ ≤ 0.1. Sequences that prefer chain-solvent interactions to intra-chain interactions will be expanded relative to the FRC limit. This leads to positive values of v_es_ and corresponds to values of Δ that are greater than 0.1.

We extended our analysis of sequence-specific effective solvation volumes to naturally occurring disordered linkers in multi-domain proteins within the non-redundant human proteome. Using a stringent set of criteria (see methods section) we identified approximately 100 linear multivalent proteins from the non-redundant human proteome (20,162 sequences) and extracted 226 unique linker regions (see methods for details). For each of the 226 linkers we performed all-atom simulations to quantify the sequence-specific values of Δ. The 226 unique linker sequences span a range of lengths (**Figure 3c**). We calculated the distribution of Δ values for all linkers using results from all-atom simulations (**Figure 3d**). This distribution shows that sequences of naturally occurring disordered linkers span the entire range of Δ values.

Of the 226 unique linker sequences, approximately 30% have negative effective solvation volumes (Δ < –0.1). Around 38 % of linkers have sequences defined by Δ values in the range – 0.1 ≤ Δ ≤ 0.1, implying that they will have near zero effective solvation volumes and are mimics of FRC linkers. Finally, 30% of linkers are characterized by Δ values greater than 0.1, which means that their effective solvation volumes are positive. The limiting form of a positive effective solvation volume linker is the self-avoiding random coil or SARC for which Δ ≈ 0.5. The key finding is that disordered linkers come in a range of sequence flavors, and 68% have a positive or near positive effective solvation volume.

**Table S1** provides requisite sequence details regarding the naturally occurring linkers, including the protein name, UniProt identifier (41), the value of Δ, and Gene Ontology (GO) annotations. The linkers are derived from multivalent proteins associated with a range of different functions. The proteins we identified were significantly enriched for RNA / DNA binding and RNA localization, as assessed by PANTHER-GO enrichment analysis (42) (*p* < 0.005). This is of particular interest, given that many micron-sized biomolecular condensates contain protein and RNA molecules (1). With this analysis in hand, our next goal was to understand how different types of linkers modulate the phase behavior of linear multivalent proteins.

For linkers with negative effective solvation volumes it follows that the linkers themselves can drive phase separation (43). These attractive linkers should be thought of as separate interaction domains that drive phase transitions and are hence distinct from regions that modulate the phase behavior encoded by interaction domains. Therefore, we focused our studies on disordered linkers with near zero or positive effective solvation volumes (v_es_ ≥ 0).

### Design of coarse-grained simulations to model the phase behavior of linear multivalent proteins

Numerical simulations of phase transitions require the inclusion of hundreds to thousands of distinct multivalent proteins and a titration of a spectrum of protein concentrations. Furthermore, phase transitions are characterized by sharp changes to a small number of key parameters such as connectivity and density, and the observation of these sharp transitions is computationally intractable with all-atom simulations. Therefore, we developed and deployed coarse-grained lattice models to study the impact of linkers on phase transitions.

Lattice models afford the advantage of a discretized conformational search space. This enables significant enhancements in computational efficiency. Key features of lattice models are the mapping of real protein architectures onto lattices and the design of an interaction model (3). The design of our simulation setup was inspired by the *in vitro* synthetic poly-SH3 and poly-PRM system studied by Li et al (6). The general framework of our lattice model has been extended to other systems including branched multivalent proteins (3), and is transferable through phenomenological or machine learning approaches (44) to any system of multivalent proteins and polynucleotides

We modeled each multivalent poly-SH3 and poly-PRM protein using a coarse-grained bead-tether model (**Figure 4**). A single lattice site was assigned to each SH3 domain. This sets the fundamental length scale in our simulations. Each PRM comprises of approximately 10-residues, thus giving it the approximate dimensions of a single SH3 domain. Therefore, each PRM was also assigned to a single lattice site. Previous all-atom simulations showed that the spatial dimensions of a single SH3 domain corresponds to ~7 linker residues, if v_es_ ≥ 0 (45). Therefore, the linker length can be written as *N* ≈ 7*n* where *n* is the number of lattice sites corresponding to a linker and *N* is the number of linker residues. All simulations were performed on 3-dimensional cubic lattices with periodic boundary conditions. Individual SH3 domains and PRMs can bind to one another and form a 1:1 complex with an intrinsic binding energy of –2*k*_*B*_*T*. Here, *k*_*B*_ is Boltzmann’s constant and *T* is temperature. This intrinsic affinity reproduces measured dissociation constants for SH3 domains and PRMs (6).

**Figure 4:**
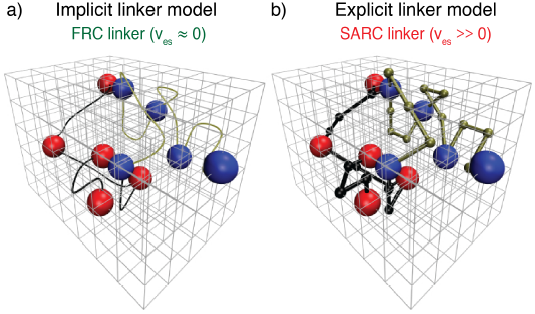
Coarse-grained bead-tether lattice models for modeling the phase behavior of multivalent proteins. All simulations were performed using 3-dimensional cubic lattice models. In these models, poly-SH3 and poly-PRM proteins were modeled as bead-tether polymers where the red beads mimic an SH3 domain, the blue beads mimic PRMs, and the black or gold tethers mimic linkers that connect domains / modules to one another. Two beads cannot occupy the same lattice site. Panel (a) shows an implicit linker model. To mimic FRC linkers, implicit linkers ensure that two tethered beads cannot move apart beyond a maximum distance, but the linker itself does not occupy any lattice sites. Panel (b) shows the explicit linker model. To mimic SARC linkers, explicit linkers consist of non-interacting beads corresponding to a prescribed number of lattice sites. The explicit linkers tether two folded domains together, but other than occupying sites on the lattice they do not engage in interactions with one another or with the interaction domains. Note that in the explicit linker model each linker bead and interaction domain occupies a single lattice site. This choice was motivated by previous analysis of the comparative effective solvation volumes of FRC and SARC linkers (45). In the figure, the linker beads are represented as being smaller than the interaction beads to emphasize that they are linkers. The real simulation box used is much larger than the lattice dimensions pictured here, which is just for illustration purposes.

We start with two stylized linkers namely, Flory random coil (FRC) linkers and the self-avoiding random coil (SARC) linkers. FRC linkers correspond to chains with v_es_ = 0. We model FRC linkers as implicit linkers (**Figure 3a**) – the linkers have a fixed length and tether the domains together, but do not occupy any volume on the lattice. Practically this is realized by imposing a cubic infinite square well potential to ensure that the lattice spacing between tethered interaction domains does not exceed *n,* the linker length in terms of the number of lattice sites. For the SARC linkers with positive v_es_, we use explicit linkers as shown in **Figure 3b**. A SARC linker of length *n* has *n* beads, where each bead is constrained to occupy vertices adjacent to its nearest neighbor beads on the lattice. Each explicitly modeled linker bead occupies a finite volume corresponding to one lattice site.

### Distinguishing between phase separation and gelation

Phase separation results from a change in density. We quantify a parameter ρ, which we define as the ratio of *R*_lattice_ to 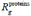 Here, *R*_lattice_ is the radius that we would obtain if all proteins were uniformly dispersed across the lattice (**Figure 5**). Conversely, 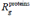 is the actual ensemble-averaged radius of gyration over the spatial dimensions of the SH3, PRM, and linker beads (**Figure 5**). For a system that has undergone phase separation, the parameter ρwill be > 1. ρ is directly related to the relative density of the proteins and measures the extent of spatial clustering of domains and linker residues. If ρ is =1, then the proteins are uniformly dispersed through the lattice.

We quantify gelation in terms of the fraction of molecules in the system that are part the single largest cluster. This is denoted as ϕ_c_ (**Figure 5**). We analyze each configuration of multivalent proteins to detect the formation of connected clusters. Within each configuration, each molecule is a *node*. An *edge* is drawn between two *nodes* if an SH3 domain from one molecule interacts with a PRM from another molecule. The connected cluster with the largest number of nodes is designated as the largest cluster and the number of molecules corresponding to this cluster is recorded. This quantity is calculated across the entire ensemble of configurations in order to generate an ensemble averaged value of ϕ_c_ for the system of interest.

**Figure 5:**
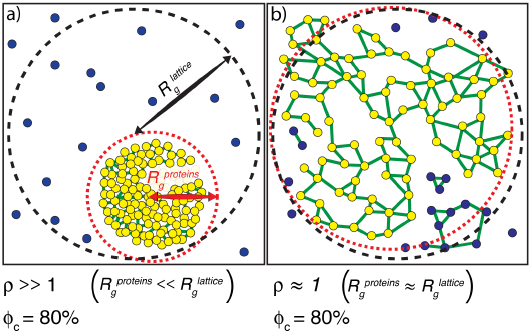
Illustration of how rand ϕ_c_ are calculated. (a) *The scenario where* ρ *>> 1*. The radius of gyration over all proteins is the root mean square distance of each of the proteins from the center of mass of the system of proteins and is depicted as the radius of the dashed red envelope. Although the red envelope is centered on the cluster, it extends beyond the cluster boundary due to the presence of proteins outside of the cluster; *i.e*., 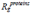 is always calculated over *all* proteins in the system. When a majority of the proteins are spatially clustered, the calculated 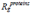 is considerably smaller than the radius of the lattice, and hence the ratio 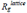 is shown as a black dashed envelope. In panel (a) a majority of the proteins are found within a single droplet-spanning cluster. This cluster encompasses ~80% of the modules, hence ϕ_c_ ~80%. Modules belonging to the single largest system spanning clusters are shown in yellow, the crosslinks are shown in green, and the “system” here refers to the droplet. (b) *The scenario where* ρ *≈ 1*. In this case, the modules are dispersed across the lattice volume as shown by the fact that the dashed red envelope is essentially coincident with the dashed black envelope. Here, we depict a scenario where 80% of the modules are incorporated into the single largest system spanning cluster, where the “system” volume corresponds to that of the entire lattice.

### Multivalent proteins with FRC linkers undergo phase separation plus gelation

We performed a series of Monte Carlo simulations using a coarse-grained lattice model for poly-SH3 and poly-PRM systems of valence 3, 5, and 7 and all combinations of these valencies. Unless otherwise specified, in all of our simulations, the linker length *n* was set to five lattice sites, approximately 35 residues. This linker length corresponds to the main mode in the distribution of linker lengths shown in **Figure 3c**.

The first row of plots in **Figure 6** shows how ϕ_c_ changes for different simulated systems and provides a quantification of gelation. Each sub-plot in **Figure 6a** shows the value of ϕ_c_ as a function of the concentrations of SH3 domains and PRMs for a particular combination of PRM and SH3 domain valence. **Figure 6a** establishes two distinctive features of multivalent systems: For a given combination of SH3 and PRM valencies, we observe a sharp increase in the values of ϕ_c_ as the concentrations of SH3 domains and PRMs increase. This behavior is consistent with the expected features of a sol-gel transition. Second, as valence increases, there is a lowering of the module concentrations at which ϕ_c_ increases sharply.

**Figure 6:**
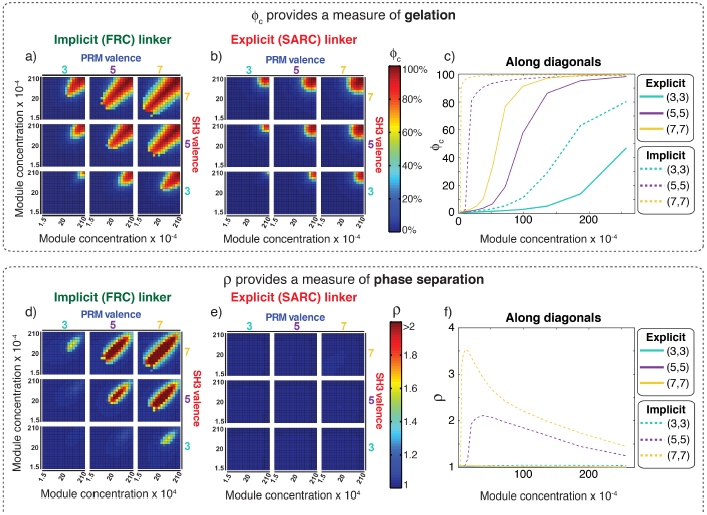
Comparative analysis of the connectivity and density transitions for multivalent proteins of fixed linker lengths. (a) Heat maps showing ϕ_c_ as a function of changes to SH3 and PRM concentrations for multivalent proteins with FRC linkers. Progression from cool to hot colors leads to the incorporation of most of the modules into the single largest cluster. The module concentrations at which sharp changes in connectivity are realized will decrease with increasing valence. (b) Heat maps equivalent to those of panel (a) for multivalent proteins with SARC linkers. (c) Analysis of how ϕ_c_ changes with module concentration for equal concentrations SH3 modules to PRMs. The solid curves plot ϕ_c_ for proteins with SARC linkers and the dashed curves are results for FRC linkers. The legend provides an annotation of the color scheme for the different curves. (d) Heat maps showing r as a function of changes to SH3 and PRM concentrations for multivalent proteins with FRC linkers. Comparison to panel (a) shows the congruence between changes to ρand ϕ_c_, especially for the 5:5, 5:7, 7:5, and 7:7 systems. (e) Heat maps. howing ρ as a function of changes to SH3 and PRM concentrations for multivalent proteins with SARC linkers. The value of ρdoes not change and remains close to one irrespective of the valence or module concentration. (f) Analysis of how ρ changes with module concentration for equal concentrations SH3 modules to PRMs. The solid curves are for proteins with SARC linkers and this shows that ρ ≈ 1, irrespective of the module concentrations. The dashed curves, for the 5:5 and 7:7 systems with FRC linkers show a sharp change above a threshold concentration of the modules. The behavior at high module concentrations is partly an artifact of our approach to increasing concentrations in the simulations, which involves fixing the number of modules and decreasing the volume of the simulation box. Accordingly, the radius of the lattice will decrease, thus decreasing ρ. However, ρ is greater than tone above a critical concentration, thus emphasizing the coupling between phase separation and gelation for proteins with FRC linkers.

**Figure 6b** shows ϕ_c_ results obtained for poly-SH3 and poly-PRM systems with SARC linkers. Here, five beads were modeled explicitly for each of the linkers between SH3 domains and PRMs. Although most systems show a sharp increase in ϕ_c_ past a threshold SH3 / PRM concentration, the concentrations at which the transitions are realized are at least an order of magnitude higher than those observed for the systems with FRC linkers. The differences between FRC and SARC linkers are summarized in **Figure 6c**, which shows how ϕ_c_ changes with module concentrations for the symmetric 3:3, 5:5, and 7:7 systems along the diagonals for equal ratios of SH3 domains and PRMs. The *x* : *y* designation refers to the *valence of SH3 domains* : *the valence of PRMs*. The value of ϕ_c_ changes sharply with concentration and this change becomes sharper as the valence increases. For a given valence, ϕ_c_ increases more sharply and at lower module concentrations for proteins with FRC as opposed to SARC linkers. This analysis shows that the effective solvation volumes of linkers can have a profound impact on sol-gel transitions.

The bottom row in **Figure 6** shows how ρchanges for each of the multivalent systems and provides a quantification of phase separation. **Figure 6d**, which summarizes the results for FRC linkers, shows sharp changes to ρ as valence increases. This recapitulates the observations in **Figure 6a** for ϕ_c_ indicating that changes to connectivity are coupled to changes in density. This is illustrated in plots for the 7:7, 7:5, 5:7, and 5:5 systems. In contrast, the 5:3, 3:5, and 3:3 systems show gelation transitions with negligible changes to ρ. In the highly asymmetric 7:3 and 3:7 systems, the changes in ρ are considerably less pronounced when compared to changes in ϕ_c_in each simulation, the initial conditions correspond to the multivalent proteins being randomly dispersed across the cubic lattice (see **movie S1**). The movie and comparative analysis of results in **Figures 6a** and **6d** provide visual support for the suggestion that systems with FRC linkers undergo phase separation plus gelation.

**Figure 6e** shows the results obtained for poly-SH3 and poly-PRM systems with SARC linkers. The results provide a striking contrast to the results obtained for proteins with FRC linkers (see **movie S2** in the supplementary material). None of the systems show discernible changes to ρ. This implies that sol-gel transitions are realized only when the concentrations are large enough to enable networking through random encounters. The positive effective solvation volumes of SARC linkers suppress phase separation and these systems undergo gelation without phase separation. **Figure 6f** summarizes the distinctions between FRC and SARC linkers by plotting ρ versus the concentration of modules for the symmetric cases with equal ratios of SH3 domains and PRMs. For SARC linkers, ρ ≈ 1 across the entire concentration range for (solid curves). This emphasizes the suppression of phase separation for systems with SARC linkers. For proteins with FRC linkers, the values of ρ increase sharply above unity beyond system-specific critical concentrations.

Representative post-equilibration configurations for 7:7 systems with FRC and SARC linkers of length five are shown in **Figure 7**. Both snapshots correspond to values of ϕ_c_ being above the gel point. The bounding box corresponds to the volume of the simulation cell and provides perspective regarding the change in density and connectivity within the system. In **Figure 7a**, a dense (high ρ) spherical droplet, which is a gel (ϕ_c_ is above the percolation threshold), coexists with a dilute sol of well-dispersed proteins. In contrast, **Figure 7b** shows how a system spanning network, *i.e*., gelation occurs in the absence of phase separation.

**Figure 7:**
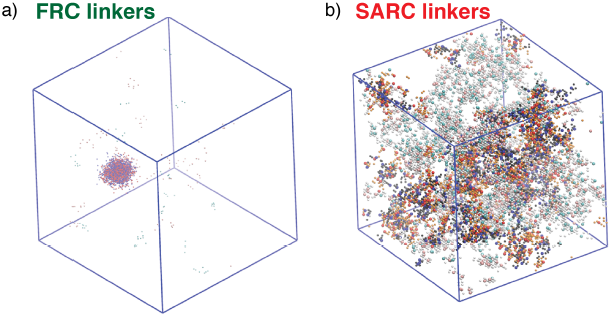
Representative, post-equilibration, snapshots for the 7:7 system above the gel points with FRC, panel (a), and SARC linkers, panel (b) of length *n* = 5. In panel (a), the SH3 modules are shown in red and the PRMs in blue. In panel (b), the coloring is similar to panel (a). Additionally, molecules that are part of the single largest, system spanning cluster are shown in orange.

### Linkers influence the degree and type of cooperativity in sol-gel transitions

If the linkers are short, then irrespective of the effective solvation volume, the formation of a physical crosslink between a pair of multivalent proteins will increase the probability that a second crosslink can form between the same pair of proteins. In this scenario, there is *positive local cooperativity,* in that the apparent affinities will increase (46) but the network cannot grow because the apparent valence is lower than the actual valence. In the limit of positive local cooperativity, phase separation and gelation are suppressed because collective interactions amongst the molecules are weakened in favor of forming network terminating dimers and oligomers. This scenario corresponds to *infinite negative global cooperativity*. In this scenario, there will neither be gelation nor phase separation plus gelation.

For long enough linkers the domains become independent of one another. Here, the extent of crosslinking and the gel point are determined entirely by the valence of domains and the intrinsic affinities between domains. This is the limit of classical Flory-Stockmayer theories with *zero local cooperativity*. The linkers are passive tethers that generate multivalency, but they do not make any other contributions to the transitions of multivalent systems. In the limit of zero local cooperativity, gelation occurs without phase separation, implying *zero global cooperativity*.

For intermediate linker lengths, the signs and magnitudes of the effective solvation volumes of linkers will determine the overall phase behavior. Disordered linkers with negative or near zero v_es_ values can enable phase transitions characterized by *positive global cooperativity* because they can drive density transitions of multivalent proteins. These linkers can be confined to small volumes, when compared to the volume of the entire system. This derives from the preference for chain-chain interactions (v_es_ < 0) or indifference for chain-chain versus chain-solvent interactions (v_es_ ≈ 0). Increased concentrations of domains within confined volumes realized by density transitions will enable connectivity transitions because the gel point is lower than the concentration of domains within the dense phase. If a multivalent protein contributes to growth of a network by forming a crosslink with a free domain on a protein that has already formed a crosslink with another protein, then the increased crosslinking enables gelation. These collective effects can also increase the apparent affinities between domains (as in the first scenario) thereby increasing the concentration of interaction domains. Increased crosslinking enables a connectivity transition whereas increased concentration of domains enables a density transition. The regime of positive global cooperativity corresponds to the regime where phase separation plus gelation is realized.

Linear multivalent proteins with large positive effective solvation volume linkers (v_es_ >> 0) will engender *negative global cooperativity* because the linkers prefer to be solvated and will resist confinement within droplets. In this sense, linkers with large positive effective solvation volumes are analogous to solubilizing tags. Additionally, due to their large positive effective solvation volumes, the linkers act as obstacles that inhibit interactions between domains. These linkers decrease the apparent affinity between interaction domains and reduce the degree of crosslinking. Accordingly, the ability to concentrate multivalent proteins is weakened, and so is the ability to grow a system-spanning network via a connectivity transition. In the scenario of negative global cooperativity, phase separation is suppressed and gelation is realized at bulk concentrations that are considerably higher than the Flory-Stockmayer limit. As a reminder, linkers do not make any contribution to determining the gel point in the Flory-Stockmayer limit (21-23, 26), only the valence and intrinsic affinities matter.

To summarize, phase separation plus gelation leads to positive global cooperativity, and enables the formation of a percolated network at bulk concentrations that are considerably smaller than the Flory-Stockmayer limit. Systems with zero or negative global cooperativity undergo gelation without phase separation and sol-gel transitions occur at or above the Flory-Stockmayer limit.

### A dimensionless parameter to quantify cooperativity

To put the ideas described above on a quantitative footing and enable comparisons across different systems we calculated the percolation threshold for ϕ_c_, which we designate as ϕ_cc_ and use this to quantify the gel point *c*_*g*_. The gel point is the concentration threshold beyond which the system crosses the percolation threshold. The methods for computing ϕ_cc_ for a system with prescribed values for the valence and the binding energy between interaction domains, as well as the calculation of the gel point from ϕ_cc,_ are described in the methods section.

We introduced a dimensionless parameter *c\** to quantify the magnitude and type of cooperativity that characterizes phase transitions of linear multivalent proteins. A measure of cooperativity also directly reveals the nature and extent of coupling between phase separation and gelation. The parameter *c*\* is defined as the ratio of *c*_*g*,sim_ to *c*_*g*,FS_. Here, *c*_*g*,sim_ is the gel point quantified in simulations with linkers of specified length and effective solvation volume. It is defined as the lowest concentration of modules at which ϕ_c_ > 0.17 (see methods section). This is percolation threshold for our system of finite-sized linear multivalent proteins (see methods section). In contrast, *c*_*g*,FS_ is the gel point obtained from Flory-Stockmayer theories (21-23, 26) (see methods section). Therefore, the value of *c*_*g*,FS_ provides an important touchstone for quantifying the influence of linkers on phase transitions, and provides a measure of the deviation from the mean-field behavior expected of long inert linkers.

The value of *c\** can be less than one, equal to one, or greater than one, depending on whether the system is characterized by positive, zero, or negative, global cooperativity, respectively. It is worth emphasizing that *c*\* quantifies the joint effects on changes to the apparent affinities of interaction modules and the extent of crosslinking. Therefore, *c*\* measures the extent and nature of coupling between phase separation and gelation. Importantly, no temporal order of operations is implied in the calculation or analysis of *c*\*.

### FRC linkers have an optimal range of lengths for positive cooperativity

We quantified the impact of linker lengths on the degree and magnitude of cooperativity for FRC linkers. **Figure 8a** shows a plot of *c\** as a function of linker lengths for 3:3, 5:5, and 7:7 systems with FRC linkers. The profile of *c*\* is non-monotonic. In the short linker limit, *n* ≤ 2, the value of *c\** is greater than one. Because these linkers are too short, complexes terminate in dimers of poly-SH3 and poly-PRM proteins. This is the regime of positive local and negative global cooperativity where phase transitions do not occur.

For multivalent proteins with a valance of 5 or 7 and linker lengths in the range 3 ≤ *n* < 12 (or 21 ≤ *N* ≤ 84, where *N* is the number of linker residues), the value of *c\** is less than one, and the lowest values of *c\** are realized for linkers of length 3 < *n* < 6. FRC linkers within a defined length range engender positive global cooperativity and for linker lengths in this optimal range, positive global cooperativity increases with increasing valence. Positive global cooperativity weakens with increasing linker lengths. Hence, for long linker lengths, *c\** converges to one implying that the domains interact independently when the FRC linkers are sufficiently long. This is the regime of zero global cooperativity.

**Figure 8:**
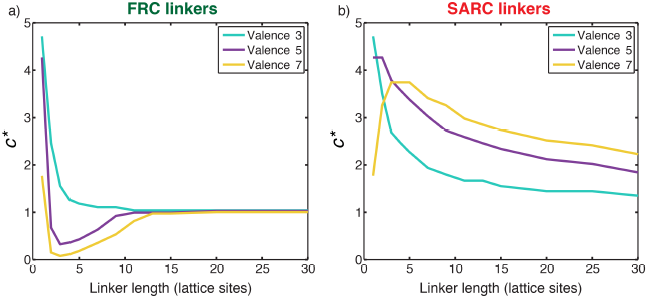
Quantifying cooperativity and the coupling between phase separation and gelation. (a) Plot of *c\** as a function of linker length for three symmetric multivalent systems connected by FRC linkers. There is an optimal range for linker lengths where *c\** < 1, implying positive global cooperativity that gives rise to phase separation plus gelation. For long linkers, *c\** converges to unity, implying an absence of cooperativity and pure sol-gel transitions, in accord with Flory-Stockmayer theories. (b) Plot of *c\** as a function of linker length for three symmetric multivalent systems connected by SARC linkers. The value of *c\** is greater than unity for all linker lengths. This points to the suppression of phase separation by linkers with positive effective solvation volumes, and a shifting of the gel point to higher concentrations compared to the Flory-Stockmayer threshold.

### SARC linkers lead to negative global cooperativity

**Figure 8b** shows a plot of *c\** as a function of linker lengths for 3:3, 5:5, and 7:7 systems with SARC linkers. Here, *c\** is greater than one for all the linker lengths. This is a signature of negative global cooperativity. Linkers with positive effective solvation volumes suppress phase separation and shift the gel point to higher concentrations when compared to the threshold predicted by Flory-Stockmayer theories. Explicit linkers also lower the apparent affinity through negative global cooperativity because their positive effective solvation volumes promote solvation thus diminishing productive associations among domains. This becomes less of an issue as the linkers become longer. If one corrects the intrinsic affinity to account for the weakened apparent affinity, then the convergence of the systems with long linkers to the Flory-Stockmayer limit is recovered (not shown). However, the profiles do not change qualitatively and this points to fundamental differences between systems with FRC versus SARC linkers.

### Phase diagrams delineate parameters for distinct types of phase transitions

**Figure 9** shows the phase diagram that we computed from concentration dependent simulations for a 5:5 system and a hybrid five-site linker. This phase diagram is shown in the two-parameter space of the concentration of domains along the abscissa and increasing intrinsic affinities along the ordinate. For affinities below 3*k*_*B*_*T*, the system undergoes a continuous transition from a sol to a gel and the green dashed line demarcates the sol-gel line. The gels correspond to system spanning networks that percolate through the entire simulation volume. The critical point for this system, shown as a red asterisk, is defined jointly by a critical interaction affinity (3*k*_*B*_*T*) and a critical module concentration (~ 10^−3^ polymers / voxel).

Above the critical point, the system undergoes phase separation plus gelation. As the interaction affinity increases above 3*k*_*B*_*T*, the system separates into two coexisting phases namely, a dilute phase, which is a sol, and a dense phase, which is a gel. As an illustration, for an interaction affinity of 4.5*k*_*B*_*T*, the coexisting concentrations that define the two phases are depicted as intercepts along the abscissa, and designated as *c*_*sl*_ and *c*_*sh*_, which are respectively the concentrations of dilute and dense phases. Notice that the gel point, *c*_g_, defined as the concentration beyond which the percolation threshold, ϕ_c_ > 0.17, lies within the two-phase regime such that *c*_*sl*_ *< c*_*g*_ < *c*_*sh*_. Here, *c*_*g*_ is the apparent gel point that is extrapolated by extending the green dashed line in **Figure 9**. Accordingly, the density transition, which we quantify as the concentration range above which ρbecomes greater than 1.08, enables gelation because the concentration within the dense phase (*c*_*sh*_) is higher than the apparent gel point (*c*_*g*_).

**Figure 9:**
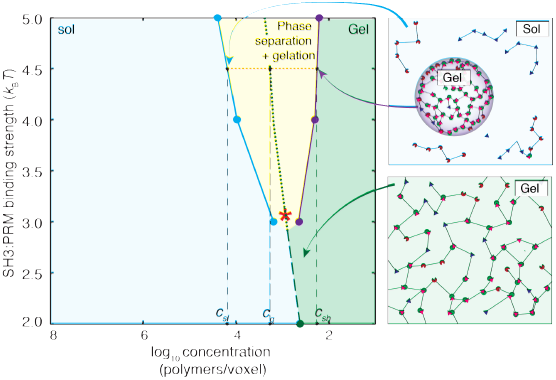
Phase diagram for a 5:5 system with a hybrid five-site linker. Here, for each linker, two of the linker beads were modeled explicitly, while the other three were modeled implicitly. For low binding affinities between SH3 domains and PRMs (< 3*k*_*B*_*T*), the system undergoes a continuous sol-gel transition as a function of module concentration, and the affinity-specific gel points lie on the green dashed line. The red asterisk denotes the critical point located at an interaction affinity of ~3*k*_*B*_*T* and a module concentration of ~10^−3^ polymers / voxel. Above an interaction affinity of ~3*k*_*B*_*T*, the system undergoes phase separation plus gelation. Phase separation is characterized by a coexistence curve with two arms, shown in blue and purple. A solution with a bulk concentration that falls within the yellow region will never form a one-phase solution. Instead, it will separate into coexisting dilute and dense phases. The concentrations within these phases are equal to the concentrations taken from coexistence curves that intersect with the corresponding tie line (red dotted line). This is illustrated for interaction strengths of 4.5*k*_*B*_*T*. Any solution with a bulk concentration along the tie line will phase separate into a dense phase and a dilute phase of a fixed concentration *c*_*sl*_ and *c*_*sh*_, respectively. For this system, the high concentration arm of the coexistence curve always lies beyond the gel-line, and therefore, the dense phase will always form a gel. The gel line within the two-phase region is calculated based on the percolation threshold and is shown as a dotted green line, which is really an extrapolation of the green dashed line. It highlights the fact that *c*_*sl*_ *< c*_*g*_ *<c*_sh_ throughout the two-phase regime. The callouts on the right show schematics of the dilute sol coexisting with a dense gel (top right) and a system spanning gel that forms via gelation without phase separation (bottom right).

The width of the two-phase regime increases with interaction affinity. This implies that phase separation is realized at lower concentrations of the interacting domains and is depicted by a leftward shift of the arm shown in light blue in **Figure 9**. Concomitantly the gel becomes more concentrated and this is depicted by a rightward shift of the arm shown in purple in **Figure 9**. Therefore, if the linker sequence is fixed, mutations to interaction domains or SLiMs that increase affinity will enhance phase separation plus gelation, giving rise to concentrated gels that coexist with dilute sols.

### Phase separation is destabilized as the effective solvation volumes of linkers increase

The effective solvation volumes of linkers were titrated by fixing the linker length and changing the number of linker beads that were modeled implicitly versus explicitly. The magnitude of the effective solvation volume is quantified in terms of the number of explicitly modeled beads within each linker. For example, if two out of five linker beads are modeled explicitly, then v_es_ is proportional to the volume of two lattice units as is the case for linkers that yield phase diagrams shown in **Figures 9 and 10c**.

Each of the panels in **Figure 10** corresponds to a distinct type of linker, defined by the effective solvation volume, *i.e*., the number of explicitly modeled linker beads for a linker of length five. Progressing from the top left corner to the bottom right corner, we find that the critical point shifts to higher interaction affinities as the effective solvation volumes of linkers increase. If the linkers have more of an FRC-like character, then there is a high likelihood that phase transitions occur via phase separation plus gelation. For a given value of the affinity, the width of the two-phase regime increases as the magnitude of the effective solvation volume decreases. In contrast, the two-phase regime becomes negligibly small as the magnitude of the linker effective solvation volume increases. In fact, for high linker effective solvation volumes, the presence of a two-phase regime is only discernible for very high affinities and phase transitions occur mainly via continuous sol-gel transitions.

**Figure 10:**
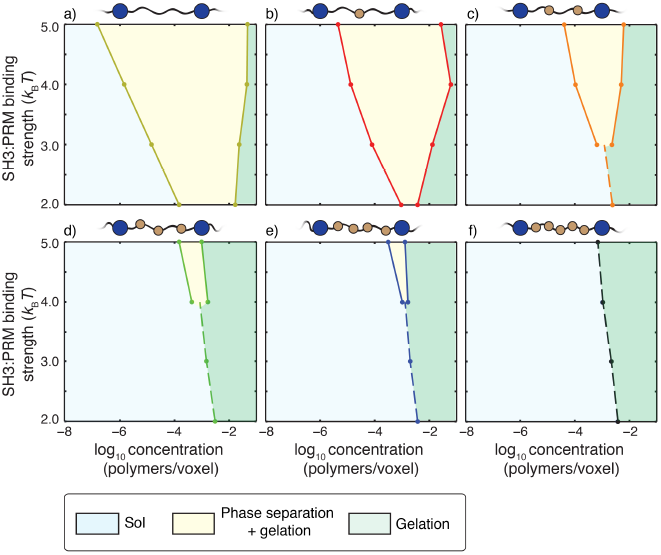
Impact of linker v_es_ values on coupling between phase separation and gelation for 5:5 systems with linkers of length *n* =5. Progressing from panel a) to panel f), the value of v_es_ for each of the linkers increases from 0 to 5 in terms of number of lattice units. The widths of the regimes that correspond to phase separation plus gelation (yellow regions) shrink as the effective solvation volumes of linkers increase. The sol-gel lines are shown as dashed lines in each panel. For a) and b) the sol-gel transitions without phase separation are realized for SH3 : PRM affinities that are weaker than 2*k*_*B*_*T* and hence they are not shown in these panels. Each panel is annotated with a schematic to show the design of hybrid linkers and each schematic we shown only a single linker for clarity.

## Discussion

Using numerical simulations, we showed that that linear multivalent proteins can undergo two distinct types of phase transitions namely, *sol-gel transitions* and *phase separation plus gelation*. We also showed that linkers between domains / motifs in linear multivalent proteins are not just passive tethers. In addition to serving as scaffolds for motifs, as has been shown before (37, 47), the physical properties of linkers such as their lengths and effective solvation volumes will directly influence the coupling between phase separation and gelation (48).

The distinction between sol-gel transitions versus phase separation plus gelation was formalized in the theoretical work of Semenov and Rubinstein (48, 49). In their mean-field “*stickers on a chain”* model, the stickers are akin to binding domains or SLiMs in multivalent proteins and the effects of linkers between stickers can be quantified in terms of their effective solvation volumes. Semenov and Rubinstein showed that for infinitely long polymers, phase separation facilitates gelation for chains with negative, near zero, or mildly positive effective solvation volumes. The coupling between phase separation and gelation is weakened by the suppression of phase separation as v_es_ becomes positive. They also showed that the coupling between phase separation and gelation is modulated by the affinities between stickers.

Our numerical results summarized in **Figures 6-10** are in accord with the theoretical predictions of Semenov and Rubinstein. This is gratifying given that we focus on finite-sized polymers, to which the simplifications of mean field theories are not transferrable. We have also shown that the effective solvation volumes of linkers are directly determined by their primary sequences (**Figure 3**). Additionally, we find that there is an optimal range of linker lengths that supports phase separation plus gelation for a given interaction affinity between domains.

We focused our simulations of phase transitions on linkers with zero or positive v_es_ values. However, as shown in **Figure 3d**, approximately 30% of linkers in the sub-proteome of linear multivalent proteins have negative v_es_ values. These linkers will be self-attractive. They can also engage in non-specific attractive interactions with interaction domains as well as other linkers of different sequence composition that have negative v_es_ values. Linkers with negative v_es_ values are best thought of as additional interaction sites. Therefore, the main effects of linkers with negative v_es_ values will be an effective shortening of the linker length enabling an increase in the effective valence through attractive interactions. These effects were illustrated in a previous study that was designed to study coexisting dense phases formed by the intrinsically disordered RGG domain of the protein Fibrillarin-1 (FIB1). There, the RGG domain of FIB1 was modeled using five explicit sticky beads thus conferring an effectively negative v_es_ value on this domain (3). Linkers with negative v_es_ values are likely to yield significantly more dense droplets when compared to linkers with near zero or positive v_es_ values. This is underscored in recent measurements of intra-droplet concentrations for disordered proteins with positive (32) versus negative v_es_ values (50). The intra-droplet concentration for the RGG domain of LAF-1 (32), which has a positive v_es_ value, is two orders of magnitude smaller than the intra-droplet concentration measured for elastin-like polypeptides (50), which have negative v_es_ values.

Interestingly, the sequences of many low complexity domains that tether RNA recognition modules in proteins such as hnRNP-A1 and FUS are characterized by negative v_es_ values. The high density within these droplets might explain why disease-associated mutations within these sequences engender apparently pathological sol-gel transitions that appear to be aided by conformational changes into beta-sheet-rich fibrils (51-56). In contrast, linkers characterized by mildly negative, zero, or mildly positive v_es_ values might form reasonably dilute droplets and functional gels that suppress pathological transitions (6, 7, 11, 15, 47). It is also possible that active processes inhibit gelation within dense droplets if gelation is refractory for biological function. Phase separation without gelation might be realizable in the presence of processes that shear physical crosslinks. Such a scenario would be an example of a so-called active liquid (57, 58) or more precisely a *non-equilibrium liquid* where energy is expended to suppress gelation that would accompany phase separation of multivalent proteins (17). Competitor molecules such as specific RNA sequences might also enable a shearing of percolated networks (14), although this has not been formally proven.

We speculate that the regulation of cell signaling by phase transitions might require phase separation plus gelation. This is evidenced by the formation of spherical droplets that is driven by specific multivalent proteins comprising of multiple interaction domains or linear motifs (2, 6, 7, 15, 59, 60). Sol-gel transitions that are decoupled from phase separation may also be useful in biology. Halfmann has recently reviewed functional scenarios where low complexity domains might undergo dynamical glass transitions that can resemble sol-gel transitions without phase separation (61). The glass transitions of the inactive bacterial cytosol and the transition to “solid-like” materials in fungi as a response to pH induced stresses are examples of sol-gel transitions on the whole cell level that do not have the characteristic hallmarks of accompanying phase separation of specific components (8, 9).

We further propose that effective scaffolding proteins for phase separation are likely to be linear multivalent proteins with linkers that have low effective solvation volumes (v_es_ ≈ 0). Proteins with linkers that have large positive v_es_ values are likely to be clients that partition into the droplets formed by the scaffolds (1). Further, the precise nature of phase transitions might be biologically tunable. For example, the effective solvation volumes of linkers in linear multivalent protein can be tuned through the synergistic action of kinases and phosphatases (60, 62). This will alter the fraction of charged residues along linkers thus enabling a coupling / decoupling between phase separation and gelation. Support for this proposal comes from the observation that the substrates for multisite phosphorylation tend to be enriched in disordered regions with positive effective solvation volumes (34, 35). Additionally, posttranscriptional processing of mRNA transcripts via alternative splicing can also be a route for making tissue-specific alterations to linker sequences. Interestingly, transcripts coding for disordered regions are preferentially targeted by tissue-specific splice factors when compared to transcripts for folded domains (63, 64).

Our inventory of linker sequences, shown in **Table S1**, combined with the analysis presented in our numerical simulations, provides a ready-made route to search for candidate linear multivalent proteins that drive phase separation plus gelation versus pure sol-gel transitions. Clearly, we need detailed experimental and theoretical characterization of phase diagrams of multivalent proteins, with special attention to the intersection of sol-gel lines and the two-phase regime. Our work opens the door to designing systems with bespoke sequence-encoded phase diagrams.

## Methods and analysis

### Design of the lattice model and interaction matrix

The interaction matrix includes the following terms: Each interaction domain (SH3 domain or PRM) or explicitly modeled linker bead has a finite v_es_ such that each lattice site may have only one domain or linker bead. All other interactions are nearest neighbor interactions such that adjacent sites *x* and *y* on the lattice are assigned an interaction energy ε_*xy*_ in units of *k*_*B*_*T*, where *k*_*B*_ is Boltzmann’s constant and T is ithe simulation temperature. We designate lattice sites occupied by SH3 domains using the letter S; sites occupied by PRMs by the letter P; and sites occupying linker beads by the letter L. In the default model, the interaction energies have the form: *u*_SS_ = *u*_PP_ = *u*_LL_ = *u*_SL_ = *u*_PL_ = 0 and *u*_SP_ = – 2*k*_*B*_*T*.

### Design of Monte Carlo moves for simulating the phase behavior of multivalent proteins

Five types of moves were deployed to evolve the system. (i) In addition to occupying adjacent lattice sites, two interacting domains are in a bound state if and only if this is specified by the interaction state of the domains. Accordingly, one of the moves randomly changes the interaction state of a domain without changing lattice positions. (ii) The torsional state of an end module that is tethered on one side is altered and a new interaction state is chosen at random. This attempts to move the module to a new location that is within tethering range of the linker, which is the maximum allowable length for the linker. If the module is an interaction domain, then this move also changes the interaction state of the domain similar to move 1. (iii) Crankshaft motions are applied to modules tethered on both sides. The module is moved to a new location that is within tethering range of all linkers that connect to the module in question. This is followed by randomly choosing a new interaction state if the module is an interaction domain. (iv) This move involves the collective translation of all modules that are part of a connected network. The latter is calculated by analyzing the list of all proteins that are connected through interacting domains. An arbitrary translation in any direction is then attempted. (v) Finally, individual chains are allowed to undergo reptation via a slithering motion of a protein by removing an end domain and its linker and appending it to the other end. The domain and linker are placed in a random position that maintains the tether ranges. After the new position has been assigned, the interaction state of the domain is randomly assigned.

### Acceptance and rejection of Monte Carlo moves

If a move results in placement of a domain or module on a site that is already occupied, then the move is rejected. For rotational, torsional, crankshaft, and reptation moves, the moves that do not lead to steric overlap with occupied sites are accepted according to a modified Metropolis criterion *viz*., min{1, *w*exp(−Δ *E*)}. Here, Δ*E* is the change in the energy of the system that results from proposed move. The energy is normalized with respect to *k*_*B*_*T*. The parameter *w* is set based on the proposed type of move. For rotational moves, *w*=1; for torsional and crankshaft moves, 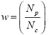, Where N_p_ and N_c_are the number of possible interacting states in the proposed and current states, respectively; finally, for reptation moves, 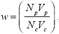, where N_p_ and N_c_ are the number of possible interacting states in the proposed and current states, respectively whereas *V*_*p*_ and *V*_*c*_ are the total number of conformations the domain and linker could be placed in the proposed state and current state respectively. These modifications to the standard Metropolis Monte Carlo acceptance criterion ensure the preservation of microscopic reversibility. The translation of a connected network does not create or destroy interactions, nor does it move the relevant linkers. Therefore, the proposed translational moves are always accepted if the move does not lead to steric overlaps.

### Production runs to generate phase diagrams

For a majority of the simulations, except those where finite size artifacts were queried or the binding affinities were titrated, the interaction energy between adjacent sites with SH3 domains and PRMs was set to –2*k*_*B*_*T*. In every system, there were 2.4 × 10^3^ interaction domains. Concentrations of domains were titrated by changing the number of lattice sites. Each simulation was run for 5 × 10^9^ steps and the average over the last half was used to calculate the size of the largest connected network.

In order to query the onset of a gelation transition, we quantified the fraction of molecules that make up the largest connected cluster within the system. We designate this as ϕ_c_ The value of ϕ_c_ that is associated with crossing the critical concentration for percolation, defined as the gel point, is determined by comparing the largest connected network from a randomly generated network to the critical concentration predicted by Flory-Stockmayer theory. Here, the number of nodes in the random network is set to the number of interaction domains used in the lattice simulations. The random network was generated for stoichiometric concentrations of complementary domains. For each domain of type A, a random number was compared to the gross probability *p* that an individual domain would be interacting with a domain of type B. If the random number was less than *p*, a partner was chosen randomly among the domains of type B that do not already have a binding partner.

### Calculating the gel points from Flory-Stockmayer theory

The gel point or more precisely, the percolation threshold for multivalent polymers can be estimated by analytical methods, one of which is based on Flory-Stockmayer theories. Here, the important parameters are the number of interacting modules within the polymers, *V*, and the fraction of bound modules, *x*. For a specific multivalent protein that is incorporated into a pre-formed network, the average number of additional proteins recruited into the network is denoted as ε and is expressed as: ε = (*V –* 1)*x*. In a system with two types of multivalent proteins *a* and *b*, such as the poly-SH3 and poly-PRM system, the average number of proteins that are recruited into a pre-formed network of multivalent proteins and their ligands can be expressed as: ε = ε_*a*_ε_*b*_ = (*V*_*a*_ *–* 1)*x*_*a*_(*V*_*b*_ *–* 1)*x*_*b*_.

If ε is greater than 1, then on average, each protein that is incorporated into the network will bring more than one additional protein with it thus expanding the network. This cascades into an infinitely large cluster of proteins. However, if ε is less than 1 then the proteins that are added are more likely to terminate the network rather than propagate it. For our synthetic poly-SH3 and poly-PRM system, we can calculate the fraction of interactions through knowledge of the dissociation constant, *K*_*d*._ We designate the SH3 domains as *a* and the PRMs as *b*. It follows that:

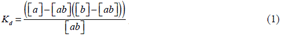

Here, [*a*], [*b*], and [*ab*] are the concentrations of SH3 domains, PRMs, and bound complexes, respectively. The concentration [*ab*] can be calculated by a simple rearrangement of equation (1), such that:

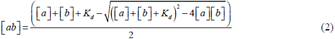

Accordingly, and

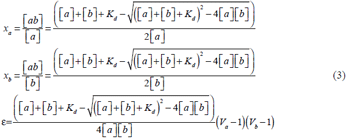

We can solve for the percolation threshold or the concentration at the gel point of module *a* as a function of the concentration of module *b* by setting ε = 1. This yields:

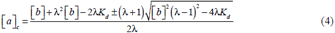

Here, λ = (*V*_*a*_ – 1)(*V*_*b*_ –1). The percolation threshold can also be calculated for the situation where [*a*] = [*b*]. In this scenario,

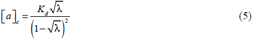

We performed simulations of random percolation models that do not account for linkers or the structure of the lattice models. Each simulation takes the valence, the number of multivalent proteins, and the fraction of bound modules as inputs. The value of ϕ_c_ is calculated for prescribed values of the fraction of bound modules and these are shown as solid sigmoidal curves in **Figure 11**. The theories of Flory (21, 22) and Stockmayer (23) can be used to calculate ϕ_cc_ analytically for given values of *V* and the binding energies, as detailed in the methods section – see equations (1) – (5). These are shown as vertical dashed lines in **Figure 11**. For a given valence *V*, the horizontal intercept that passes through intersection of the vertical dashed lines and the solid curve defines the value of ϕ_cc_. We find this value to be ≈ 0.17, irrespective of the valence. The concentration of modules at which ϕ_c_ becomes greater than 0.17 is taken to be the value of the gel point *c*_*g*_ for the system of interest. We can calculate the value of *c*_*g*_ directly from our simulations for the multivalent proteins and compare this to the value of *c*_*g*_ that is estimated from Flory-Stockmayer theories.

**Figure 11:**
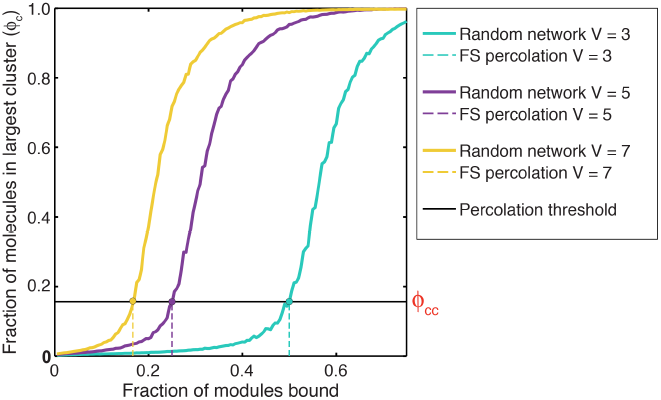
Estimating ϕ_cc_ – the critical value of the fraction of molecules in the largest cluster, ϕ_c_ that defines the gel point: To estimate ϕ**_cc,_** we plot ϕ**_c_** against the fraction of SH3 domains and PRMs that are bound. ϕ_c_ was calculated using a random network model (see methods) and for a prescribed affinity between interaction domains. ϕ_c_ shows a sigmoidal transition that shifts to the right for systems of lower valence (*V*). For each system, the dashed vertical lines quantify the percolation thresholds, which refer to the fraction of modules for a given valence *V* that must be bound in order to make a percolated network as prescribed by the theories of Flory and Stockmayer. For a given system of multivalent proteins, the intersection between the solid sigmoidal curve and the dashed vertical line quantifies the value of ϕ_cc_.

### Calculation of Phase Boundaries

We utilized r as the order parameter for differentiating between the sol-gel transitions and phase separation. The coexisting concentrations corresponding to the polymer-rich and polymer-poor phases that delineate the two-phase boundary for a given intrinsic affinity between interaction domains were calculated by assuming that the polymer-rich phase is a uniform density sphere and the polymer-poor phase has a uniform density across the remainder of the lattice. The radius of the polymer-rich phase is the radius of the sphere that is the physically relevant root of the equation:

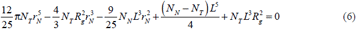

Here, *N*_*T*_ is the total number of proteins in the simulation, *N*_*N*_ is the number of proteins within the largest network, *L* is the lattice length on a side, *R*_*g*_ is the radius of gyration over all the proteins in the simulation, and *r*_*N*_ is the desired radius of the polymer-rich phase. This equation typically admits only one real root that fits within the lattice and this is true for all of our simulations. The phase boundaries were calculated using:

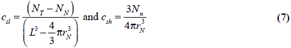

### The impact of finite sampling

In addition to starting simulations in the random coil state, we also calculated phase diagrams using simulations that were initialized from a dense phase separated state. For each simulation we equilibrated the proteins in the gel state in a box size of 34 lattice units for 5×10^9^ steps. The resulting conformation was then used to initialize simulations in a larger box by expanding the lattice boundary to achieve the desired concentration. For proteins that span the periodic boundary, the first domain was used as the reference for picking which protein image to keep. These initial conditions reproduced the critical concentrations as a function of valence and length

### All atom simulations

We identified 226 disordered linkers in the human proteome associated with multi-domain proteins. Specifically, we defined disordered linkers in multi-domain proteins as regions predicted to be disordered (65) that connected two Pfam domains (41) that were predicted or known to be folded. We then filtered for linkers that were between 15 and 200 residues in length, and sub-selected for individual proteins where two or more linkers were found. For each of these sequences all-atom simulations were run to provide a general picture of the global conformational behavior associated with disordered linkers in the human proteome.

In addition the set of disordered linkers, we also examined fourteen specifically selected sequences, each consisting of 40 residues. These sequences were chosen to enable a titration of conformational properties as a function of the sequence-encoded fraction of charged residues. Sequences of varying charge were extracted randomly from disordered regions in the human proteome. Disordered regions were identified by extracting sequences from the human proteome that were predicted to be disordered by at least five different disorder predictors in the D2P2 database. We required that each stretch have at least 40 consecutive residues that are disordered. We calculated the fraction of residues by tallying the number of ARG, LYS, ASP, and GLU residues in each fragment.

For all sequences described we performed atomistic Monte Carlo simulations using the ABSINTH implicit solvation models and forcefield paradigm (40). In this approach, polypeptide chains and solution ions are modeled in atomic detail and the surrounding solvent is modeled using an implicit solvation model that accounts for dielectric inhomogeneities and conformation– specific changes to the free energies of solvation. The simulations were performed and analyzed using tools in the CAMPARI modeling suite (http://campari.sourceforge.net). Forcefield parameters were taken from the abs_opls_3.2.prm parameter set. For each of the fourteen sequences, we performed ten independent simulations, each initialized from a distinct self– avoiding conformation. The methods used to evolve the systems and analyze the simulation results are identical to protocols used in previous studies (30, 35, 37, 66). For simulations of the 226 disordered linkers, five independent simulations per sequence were performed. Each simulation started from a distinct, randomly selected non–;overlapping conformation and comprising 5×10^6^ equilibration steps and 5×10^6^ production steps in 5 mM NaCl. Simulations of the fourteen specifically selected sequences were run for longer to obtain higher resolution statistics.

## Appendix A: Formal definition of v_es_

We start with the effective, solvent-mediated potential of mean force, which we denote as *W*(*r*). This is the free energy change associated with bringing a pair of linker residues from a non-interacting reference point to a distance *r* of one another in an aqueous solvent. Therefore, *W*(*r*) quantifies the balance of residue-solvent, solvent-solvent, and residue-residue interactions. If the residues “like” one another more than they “like” the solvent, then the effective inter-residue interactions will be attractive. If the residues “like” the solvent more than they “like” one another, then the effective inter–residue interactions will be repulsive (31).

The probability that a pair of linker residues will be a distance *r* from one another is proportional to the Boltzmann weight exp[–β*W*(*r*)], where β = (*RT*)^−1^, *T* is the temperature and R is the ideal gas constant. Because residues cannot sterically overlap with one another, the Boltzmann weight is zero for short inter–residue distances. The Boltzmann weight is one for large separations where the inter-residue interactions are effectively zero. Between these two limits, the Boltzmann weight can be large and positive for separations *r* where the inter-residue interactions are attractive. Conversely, the Boltzmann is negligibly small at inter-residue separations *r* where the effective interactions are repulsive.

The effective solvation volume per each pair of residues is defined as the negative of a integral of a function *f*(*r*) (31, 49) over the volume available to the pair of residues. Here, *f*(*r*) = exp[–β*W*(*r*)] – 1 and the integral is performed over all pairs of inter-residue separations. Depending on the inter–residue separation *r* and the type of interactions, the *f–*function will be negative (short-range steric overlaps or effective inter–residue repulsions), positive (effective inter–residue attractions), or zero (large separations). The function *f*(*r*) is known as the Mayer *f*– function and the effective solvation volume v_es_ is defined as the negative of the integral of the Mayer *f–*function over the entire volume occupied by the pair of interacting units:

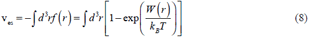

The Mayer *f*–function is a dimensionless parameter and the integral in equation (10) has units of volume. It quantifies the two–body or the effective pairwise inter–residue interactions for the polymers in solution. In terms of a virial expansion, at low concentrations, the free energy per unit volume of a polymer solution is written in terms of the polymer concentration as:

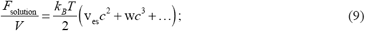

Here, v_es_ has units of volume, and w the three–body interaction coefficient, has units of (volume)^2^ and so on. In dilute concentrations where pairwise interactions dominate, which is the case when v_es_ ≥ 0, it follows that:

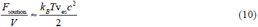

The effective interaction energy between residues is negative, zero, or positive depending on the sign of v_es_.

## Acknowledgments

We thank Salman Banani, Jeong–Mo Choi, and Kiersten Ruff for many helpful discussions. We are immensely grateful to Jill Bouchard, Cliff Brangwynne, Allan Drummond, Amy Gladfelter, Randal Halfmann, Tanja Mittag, Andrea Putnam, and Geraldine Seydoux for critical reading of the manuscript and providing us with several thoughtful suggestions that we have tried incorporate in the hope of improving the accessibility of our narrative. Grants from the National Science Foundation (MCB1614766 to RVP), the St. Jude Children’s Research Collaborative (RVP), the National Institutes of Health (RO1–GM56322 to MKR) and the Howard Hughes Medical Institute (MKR) supported this work. TSH was a graduate student scholar of the Center for Biological Systems Engineering at Washington University in St. Louis.

